# Microbial source tracking using metagenomic analysis from Cochin Estuary

**DOI:** 10.1101/2025.03.14.643239

**Authors:** Parvathi Ammini, Jasna Vijayan, Michela Catena, Deepak George Pazhayamadom, Nikhil Phadke, M Nair

## Abstract

Microbial source tracking (MST) and metagenomics offer powerful insights into microbial ecosystems and pollution sources. This pilot study used metagenomics to identify microbial pollution sources in the Cochin Estuary (CE), a vital coastal ecosystem in southwest India. The CE faces significant microbial pollution, necessitating effective tracking for conservation. Bacterial community analysis via high-throughput sequencing was conducted to determine fecal pollution sources in CE waters. Samples included human, cow, dog, and goat feces, along with sewage, slaughterhouse, and domestic waste, alongside 24 water samples from eight sites across monsoon and non-monsoon seasons. DNA extraction and 16S rRNA (V3–V4) sequencing via Illumina MiSeq enabled MST analysis. Results indicated that sewage and domestic waste were the predominant contaminants, contributing to 50% of bacterial communities. Downstream sites were more polluted than upstream areas. Proteobacteria dominated estuarine waters, whereas Firmicutes and Bacteroidetes were prevalent in source samples. This study highlights the need for targeted remediation to protect the CE’s environmental health.

**Importance of the study:** This study is essential for addressing microbial pollution in the Cochin Estuary (CE), a crucial Ramsar site facing contamination threats. Beyond identifying pollution sources, it provides valuable insights into seasonal variations in microbial composition, highlighting differences between monsoon (wet) and non-monsoon (dry) periods. It emphasizes the role of environmental factors such as conductivity, temperature, dissolved oxygen (DO), and salinity in shaping microbial communities. The study also employs advanced statistical tools like non-metric multidimensional scaling (NMDS) and principal coordinates analysis (PCoA) to assess microbial diversity and pollution patterns. Additionally, it underscores the significance of next-generation sequencing (NGS) in high-resolution microbial source tracking (MST). By integrating environmental parameters and microbial analysis, the study offers a comprehensive approach for monitoring estuarine health. These findings are crucial for policymakers, aiding in the development of targeted wastewater management and pollution mitigation strategies to protect CE’s biodiversity and ecological balance.

## Introduction

Microbial pollution is a critical threat to environmental and public health, with severe implications, particularly in low- and middle-income countries. In these regions, the impact is amplified by inadequate sanitation infrastructure, poor water management practices, and limited access to clean water. This leads to a higher prevalence of waterborne diseases like typhoid and cholera, significantly hindering socio-economic development. Identifying specific sources of microbial pollution – in particular, differentiating between human and animal origins – is essential to facilitate water quality maintenance, public health risk assessment, and environmental pollution management (1, 2).

Microbial source tracking (MST) is an innovative technique to detect microbial contamination in human-frequented water bodies (3). It comprises a suite of microbial and molecular methods used to trace fecal water pollution to its source, including metagenomic-based methods. The underlying logic behind MST is that some pathogenic bacteria are commonly found in the guts of certain hosts, and the presence of these bacteria in environmental water indicates the fecal contamination of that specific host (4). Distinguishing between human and animal fecal bacteria is important since bacteria from human feces are known to be highly pathogenic to humans. Unlike traditional water quality monitoring, MST targets specific contamination sources (5). With knowledge of the dominant sources, especially in the case of non-point source pollution, bacterial contamination can be properly addressed and halted. Determining the source of pollution provides valuable information to public health officials and environmental managers, allowing them to properly allocate tools, resources, and energy to manage better both the water and the pollution to improve water quality and public health (6–9).

Various methods, including metagenomic-based methods, have been developed to accurately track microbial sources. MST is instrumental in calculating maximum daily load allocations, assessing recreational water quality, and evaluating associated health risks. Distinguishing between human and animal fecal bacteria is crucial since bacteria from human feces are known to be highly pathogenic to humans. By addressing specific contamination sources, MST provides critical insights for public health improvement and helps prevent waterborne illnesses (7–9). Microbial pollution poses a significant threat to environmental and public health with profound implications especially in developing countries. The impact of microbial pollution in these regions is exacerbated by factors such as inadequate sanitation infrastructure, poor water management practices, and limited access to clean water sources. This leads to a higher prevalence of waterborne diseases such as typhoid and cholera and has also hindered socio-economic development. Identifying pollution sources is crucial for maintaining water quality and managing environmental pollution (1,2). Microbial source tracking (MST) plays a crucial role in identifying fecal solution sources which is management and improving water quality. MST differs from traditional water quality monitoring by targeting specific sources of contamination, providing valuable insights for public health officials and environmental managers (5). Various methods such as metagenomics have been developed to track microbial sources accurately. MST helps evaluate maximum daily load allocations, recreational water quality, and associated health risks. Distinguishing between human and animal fecal bacteria is crucial, as human waste is known to contain pathogenic microbes. MST provides valuable insights for public health improvement by addressing specific contamination sources and preventing waterborne illnesses (7–9).

This study aims to combat bacterial contamination in the Cochin Estuary (CE), home to the second largest Ramsar site in India, by identifying its sources. MST is especially important in the CE since traditional source identification is challenging due to its tidal mixing processes combined with terrigenous riverine inputs and autochthonous biological activity. The spatiotemporal dynamics of microbial communities in the estuarine environment may vitally impact water quality conditions with direct implications for human health. The CE’s significance in social, economic, and ecological terms, along with environmental stress from tourism and dense human populations, make it an ideal focus for such investigations. Previous studies have detected high levels of fecal coliforms, *E. coli*, and other pathogens (e.g., *Vibrio* and *Salmonella*) in the Periyar River, its tributaries, and in local groundwater samples (10–12), emphasizing the urgency of addressing waterborne diseases in the region. Studying microbial source tracking in the CE is particularly important due to the significant impact of terrigenous riverine inputs and the influence of autochthonous biological activity combined with tidal mixing processes. The spatiotemporal dynamics of microbial communities in the estuarine environment serve as a vital indication of water quality conditions with direct implications for human health.

Microbial community analysis utilizing next-generation sequencing (NGS) offers advantages over single marker detection by characterizing entire microbial communities simultaneously. This method distinguishes between different sources, even in regions lacking specific genetic markers or where diets differ from those in the USA and Europe (13,14). Worldwide, only a few studies have investigated the use of NGS-based community analysis for MST (13, 15–24) most of which employ 454 pyrosequencing or qPCR analysis. Presently, few studies (13, 21, 22, 24) have used Illumina MiSeq to sequence the 16S rRNA gene to characterize microbial communities in fecal and water samples for MST.

This study addresses the following questions: (1) what are the specific sources of fecal bacterial contamination in the CE? (2) What effect do the monsoon and non-monsoon seasons have on the bacterial communities and the spread of pollution from various sources? By utilizing advanced methods like metagenomics, this study identifies the spread of microbial sources of pollution in the CE to enable targeted interventions to mitigate contamination and protect the ecological balance of the estuarine system.

## Results

### Environmental Parameters

CE receives approximately 60–70% of its annual rainfall during the southwest monsoon season, and the average annual freshwater input from the CE to the Arabian sea is 22,000×10^6^ m^3^ (Rehitha et al., 2019). This intricate estuarine system experiences distinct changes due to the monsoonal cycles resulting in a 1) pre-monsoon (February–May, designated as “DP I” in this study) season when the tidal influence on the estuary is more pronounced, 2) monsoon (June–September, designated as “Wet period” or “WP” in this study) season when the riverine freshwater influx is more pronounced, and 3) a post-monsoon (October–January, designated as “DP II” in this study) season, which is an intermediate period of reduced freshwater inputs and increased tidal action. During the current study, the surface water temperature varied within a limited range (27 °C to 32.8 °C). It was lowest during the monsoon (27 °C at Station 6) and highest during the post-monsoon (32.8 °C at Station 4; **Table. S1**). Owing to the regular pre-monsoon rainfall, site Station 3, located at the confluence of the upper estuary and the Periyar River, had low salinity during this period. During the pre-monsoon, high salinity levels were primarily limited to the vicinity of both sea inlet stations (Munambam in the north and Kochi inlet located in the middle of the extent of the estuary). There were notable differences in salinity across different regions in both the dry and wet seasons. The maximum variation in salinity occurred in DP I (13.3±9.14 ppt), followed by DP II (11.1±7.29 ppt), and WP (3± 5.53 ppt). Station 4, located at the Kochi inlet, had the highest salinity during the dry season (26-28 ppt) compared to the wet season (∼16-18 ppt). During both seasons, salinity in the CE gradually dropped (0–2 ppt) in an eastward direction from the Arabian Sea towards the inland reaches. Stations 1, 2, 7, and 8 had freshwater conditions, whereas Stations 3, 5, and 6 experienced mesohaline salinities. Spatial variation in major nutrients was more pronounced than temporal variation in the CE. The pH of the samples varied from 6.50 to 7.96, while the dissolved oxygen (DO) ranged from 4.88 to 6.72 mg L^−1^ (**Table. S1**).

### Bacterial community structure in water and source samples

Water samples from eight different locations in the CE during three distinct time periods (WP, DP I, DP II) and source samples (fecal samples, raw sewage, domestic and slaughterhouse waste) were collected during 2015 and 2016. Sequencing of the V3-V4 regions of the 16S rRNA resulted in 252339.5 ± 49192.3 (mean ± SD) gene sequence reads of the eight water samples in the wet period, 263222.6 ± 71951.8 sequence reads of the eight water samples in DP II, and 164345.6 ± 32746.27 sequence reads of the eight water samples in DP I. 2530.1±53853.34 sequence reads of the source samples were generated. Based on sequencing analysis, Good’s Average was high (0.99). The percent reads to genus level ranged from 37.09% to 91.54% among all samples.

The source samples had, on average, a Shannon Index of 2.41±0.54, while the Shannon Index values of the water samples were 2.55±0.19 in WP, 2.82±0.16 in DP I, and 2.41±0.18 in DP II. Shannon diversity index values were highest at S4 during DP I and the lowest at S6 during DP II. The number of species identified was highest in DP I (1457.88 ± 171.9) followed by WP (1172.25 ± 210.2) and then DP II (929 ± 149.5) (**Table. 1**). The total bacterial diversity with more than > 95% of OTUs in the water and biological samples accounted for 32 distinct phyla.

**Table 1.** Summary of the Illumina sequencing with the number of raw and valid reads, where S1-S8 denote the sampling Stations, WP is wet period (June-Sept), DP I is dry period I (Feb-May), and DP II is dry period II (Oct-Jan).

| **Samples** | **No. of reads** | **% Reads**<br>**classifieds**<br>**to Genus** | **Shannon**<br>**Species**<br>**Diversity** | **Number of Species**<br>**identified** |
| --- | --- | --- | --- | --- |
| **Source sample** |  |  |  |  |
| Dog Fec | 211892 | 72.93% | 2.962 | 801 |
| Goat Fec | 155239 | 81.91% | 2.44 | 766 |
| Human Fec | 90697 | 72.31% | 2.782 | 601 |
| Cow Fec | 189051 | 70.22% | 2.111 | 578 |
| Sewage | 198000 | 59.09% | 3.095 | 1365 |
| Domestic Waste | 99024 | 37.09% | 2.442 | 1031 |
| Slaughterhouse | 263808 | 91.54% | 1.392 | 666 |
| **Water sample -WP** |  |  |  |  |
| S1 | 304436 | 75.65% | 2.898 | 1421 |
| S2 | 322227 | 64.49% | 2.446 | 1324 |
| S3 | 281047 | 65.69% | 2.34 | 1215 |
| S4 | 179025 | 66.01% | 2.718 | 1121 |
| S5 | 252310 | 64.20% | 2.599 | 1225 |
| S6 | 275375 | 65.91% | 2.663 | 1329 |
| S7 | 236879 | 66.95% | 2.336 | 948 |
| S8 | 167417 | 64.57% | 2.408 | 795 |
| **Water sample -DP I** |  |  |  |  |
| S1 | 274745 | 70.93% | 2.556 | 1543 |
| S2 | 367764 | 67.31% | 2.792 | 1651 |
| S3 | 173542 | 64.60% | 2.69 | 1203 |
| S4 | 237272 | 71.13% | 3.065 | 1462 |
| S5 | 260109 | 71.13% | 2.992 | 1575 |
| S6 | 323762 | 66.62% | 2.971 | 1483 |
| S7 | 337895 | 66.74% | 2.779 | 1558 |
| S8 | 130689 | 70.79% | 2.716 | 1188 |
| **Water sample -DP II** |  |  |  |  |
| S1 | 171453 | 66.93% | 2.825 | 1078 |
| S2 | 165909 | 52.63% | 2.481 | 954 |
| S3 | 127522 | 57.53% | 2.239 | 1036 |
| S4 | 189601 | 59.06% | 2.417 | 1025 |
| S5 | 210901 | 57.34% | 2.41 | 1038 |
| S6 | 176353 | 62.86% | 2.192 | 829 |
| S7 | 93251 | 63.19% | 2.36 | 827 |
| S8 | 179775 | 85.98% | 2.376 | 645 |

### Microbial community composition in the source samples

The bacterial source samples contained distinct bacterial communities that differed in the presence and abundance of certain species. Analyzing the unique bacterial profiles present in different samples enabled the detection of distinct sources. All source samples contained relatively high numbers of Firmicutes (17-61%) and Bacteroidetes (7-55%), while Proteobacteria ranged from 7-35% only. Actinobacteria (1.5-6.03%) and Cyanobacteria (0.1-1.18%) made up small fractions of the communities in all source samples. Other phyla present in the source samples were Verrucomicrobia, Fusobacteria, Spirochaetes, Tenericutes, Chloroflexi, and Nitrospirae. Among the sewage samples, Proteobacteria (36%) was the dominant phylum, followed by Bacteroidetes (28-36%) and Firmicutes (17%). The human fecal samples showed high relative abundances of Bacteroidetes (56%).

In all source samples, the prevalent classes were Clostridia, Bacteroidia, Bacilli, and Sphingobacteriia. The Clostridia were dominant in all the source samples except human feces. In sewage samples, the most dominant class was Sphingobacteriia (11%), followed by Gammaproteobacteria (10%), and Betaproteobacteria (10%). In human feces, the dominant class was Bacteroidia (54%), followed by Clostridia (29%), Gammaproteobacteria (3%), and Actinobacteria (3%) (**Fig. S1**). In dog feces, Clostridia (29%), Bacilli (22%), and Erysipelotrichi (12%) were most dominant. In domestic waste, Clostridia (44%) was followed by Flavobacteriia (10%). The dominant families in the animal fecal samples were *Clostridiaceae, Bacteroidaceae, Lachnospiraceae Flavobacteriaceae, Prevotellaceae,* and *Ruminococcaceae*, comprising over 40% of the sequencing reads from goat, dog, cow, and human feces (**Fig. 3**). In contrast, families such as *Clostridiaceae, Moraxellaceae*, and *Flavobacteriaceae* were prevalent in slaughterhouse and domestic waste samples.

**Fig. 1.**
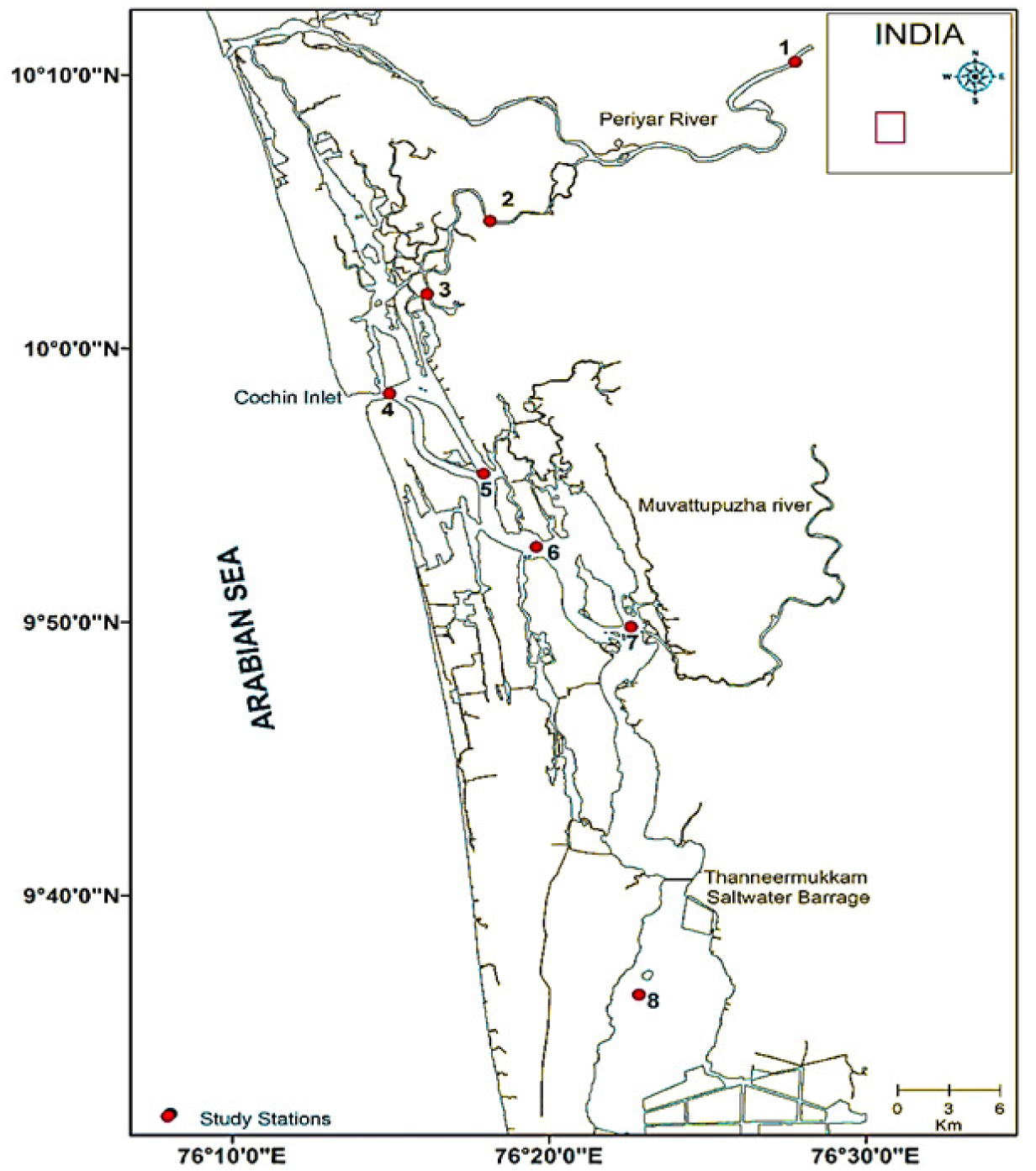
Map showing the locations of 8 sampling stations in the CE. Station 4 is the inlet Station, 1-3 are the northern Stations and 4-8 are the southern Stations.

**Fig. 2.**
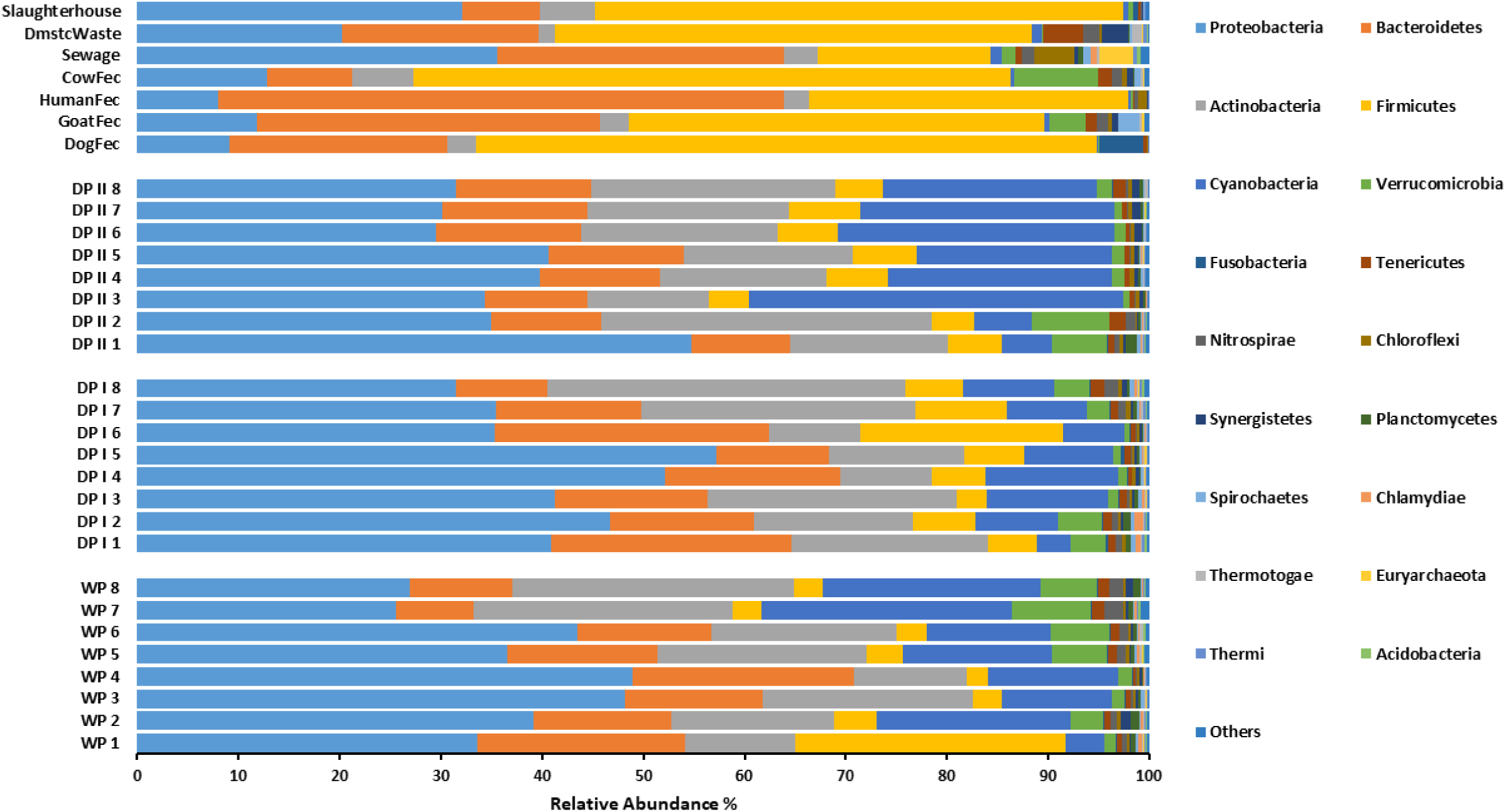
Phylum level classification at the sampling locations, where 1-8 denote the sampling Stations, WP is wet period (June-Sept), DP I is dry period I (Feb-May), and DP II is dry period II (Oct-Jan).

**Fig. 3.**
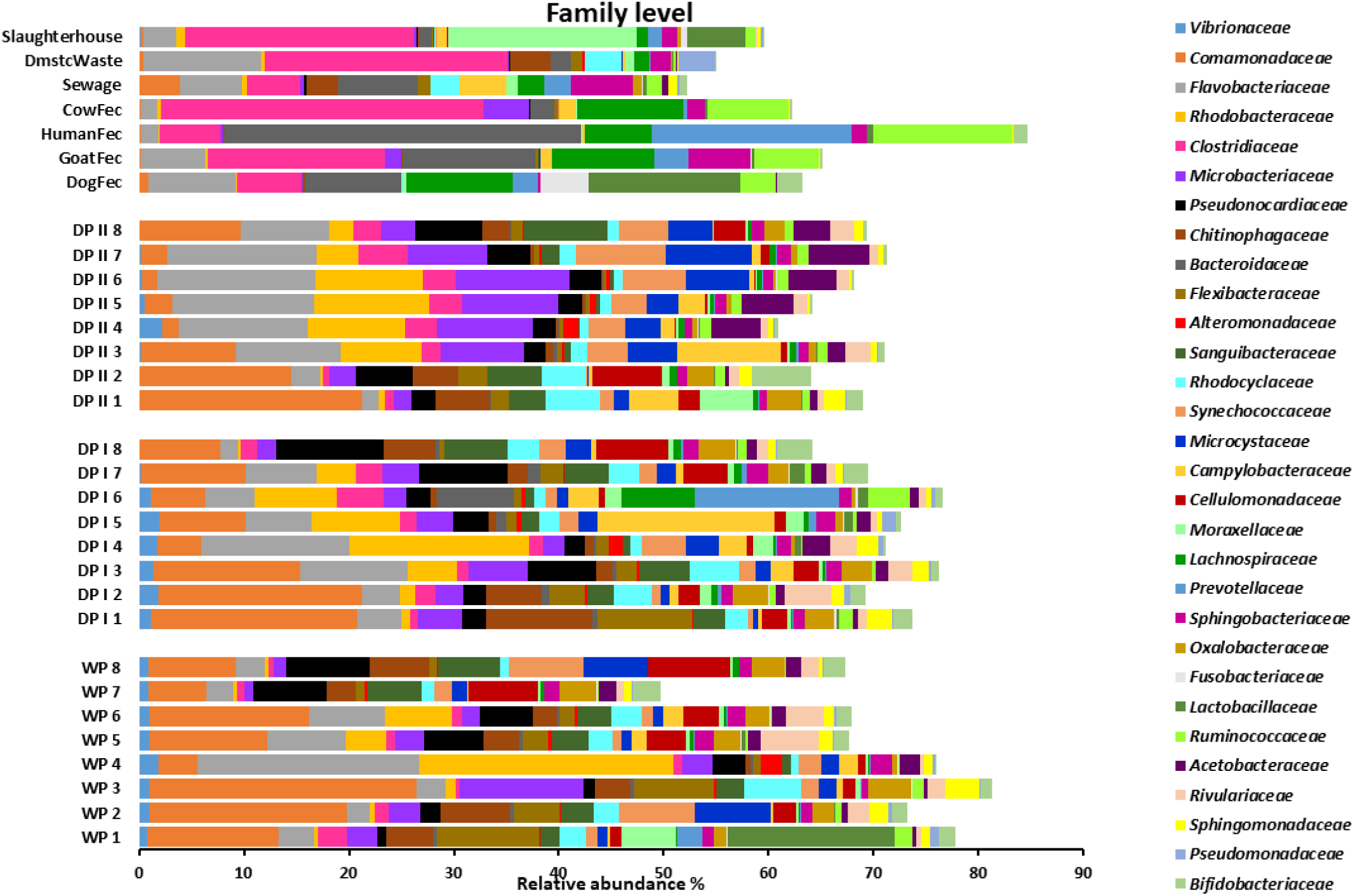
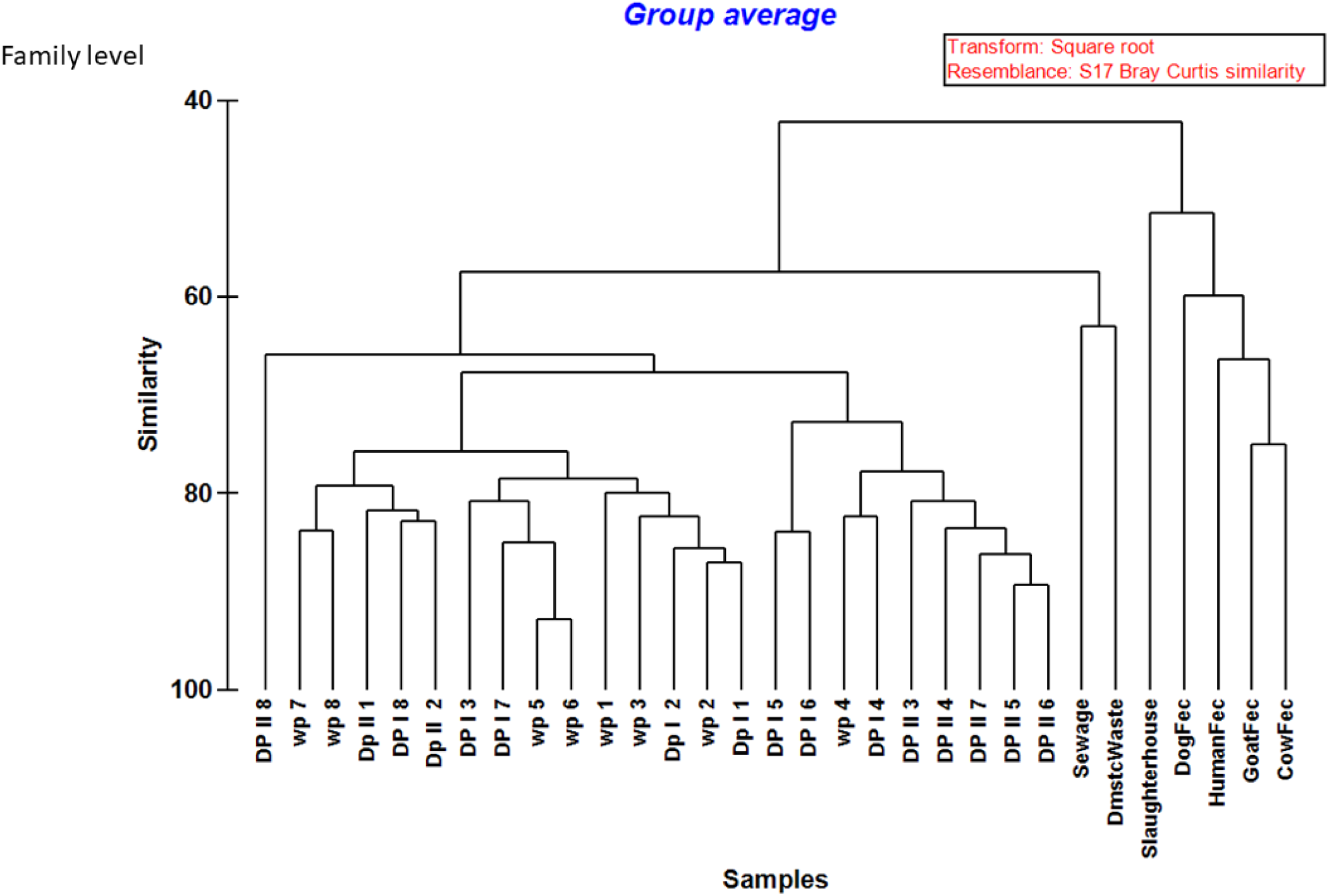
Family level distribution (top 30 families) of bacterial communities, where 1-8 denote the sampling Stations, WP is wet period (June-Sept), DP I is dry period I (Feb-May), and DP II is dry period II (Oct-Jan).

### Microbial community composition in the estuarine samples

Proteobacteria, Actinobacteria, Bacteroidetes, Cyanobacteria, Firmicutes, and Verrucomicrobiota comprised large fractions of the estuarine sample sequence reads. As expected, the estuarine metagenomes were relatively more complex compared to source samples and revealed higher diversity in all three seasons. Overall, there was a high abundance of Proteobacteria in the estuarine water samples ranging from 29-54% over the three seasons. In the wet period, Proteobacteria comprised 25-48% of sequence reads. At all Stations except 1, 2, and 4, Actinobacteria followed with 18-27% of the sequence reads. 26% of reads at Station 1 during the wet period were composed of Firmicutes, 19% were Cyanobacteria at Station 2, and 22% were Bacteroidetes at Station 4. In DP 1, Proteobacteria (31-57%) were dominant except in S8, followed by Bacteroidetes (17.47-27.12%) in Stations 1, 4, and 6 and Actinobacteria (17-61%) in the remaining Stations (2, 3, 5, 7). In DP II, there was a relatively high proportion of Cyanobacteria (19-36%) in the water samples at all Stations except 1 and 2, where Actinobacteria (15% and 36%, respectively) was dominant (**Fig. 2**).

Proteobacteria was constituted by Alphaproteobacteria, Gammaproteobacteria, and Betaproteobacteria, whereas phylum Actinobacteria was dominated by class Actinobacteria. Betaproteobacteria were more prevalent in the freshwater region of the estuary (Stations 1, 2, 3, 8) making up 5.36-35.1%, while Alphaproteobacteria were abundant in the meso and euryhaline regions (Stations 4, 5, 6, 7) comprising 8.87-28.10% (Fig S1) At the family level, the most frequent microbiota observed in environmental water communities were *Comamonadaceae, Flavobacteriaceae, Microbacteriaceae,* and *Pseudonocardiaceae.* The *Comamonadaceae* were higher in freshwater and mesohaline stations compared to the euryhaline station, especially in the dry periods, whereas *Flavobacteriaceae* were higher in mesohaline and euryhaline stations. The *Synechococcaceae* were high in DP II compared to WP and DP I (**Fig. 3**).

### Statistical analysis

Based on hierarchical clustering analysis, the animal fecal samples and slaughterhouse waste samples clustered separately from estuarine samples, but sewage and domestic waste samples clustered near the estuarine samples, indicating that the microbial community in feces varied from that found in estuarine water while the microbial communities of sewage and domestic waste were comparatively more similar to estuarine samples (**Fig. 4**).

**Fig. 4.**
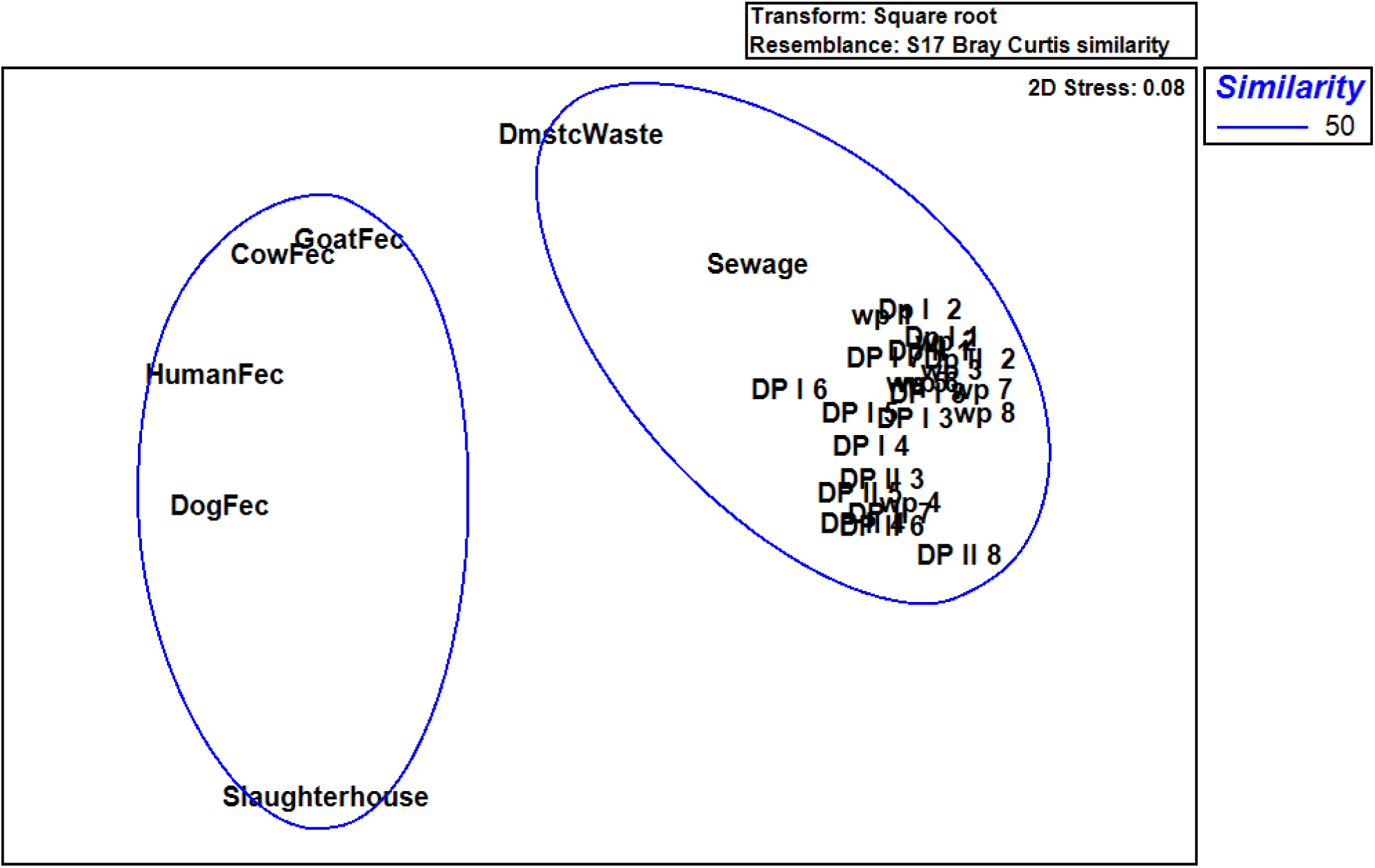
Cluster analysis of source and estuarine samples, where 1-8 denote the sampling Stations, WP is wet period (June-Sept), DP I is dry period I (Feb-May), and DP II is dry period II (Oct-Jan).

The NMDS plot illustrated the separation of samples based upon differences in microbial community structure. Based on NMDS, the environmental water samples clustered together while the animal fecal samples clustered together. Slaughterhouse waste appeared dissimilar to all other samples. Sewage and domestic waste samples clustered near to the group of clustered estuarine samples, with sewage being the closest. This could indicate the presence of sewage and domestic waste in estuarine waters (**Fig. 5**).

**Fig. 5.**
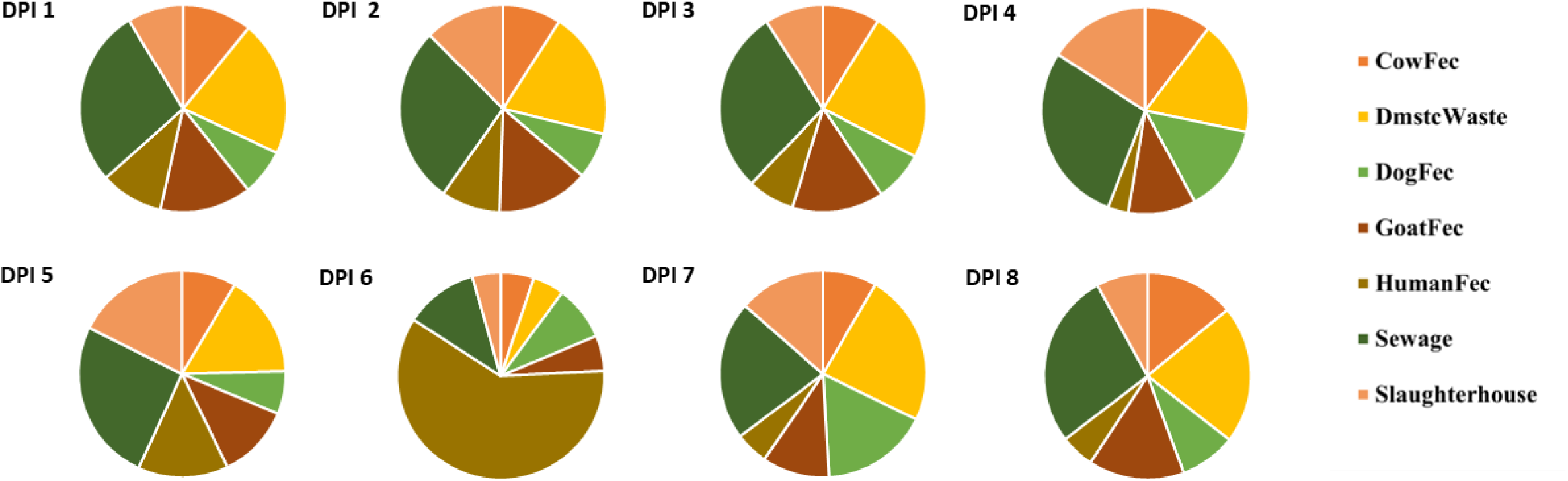
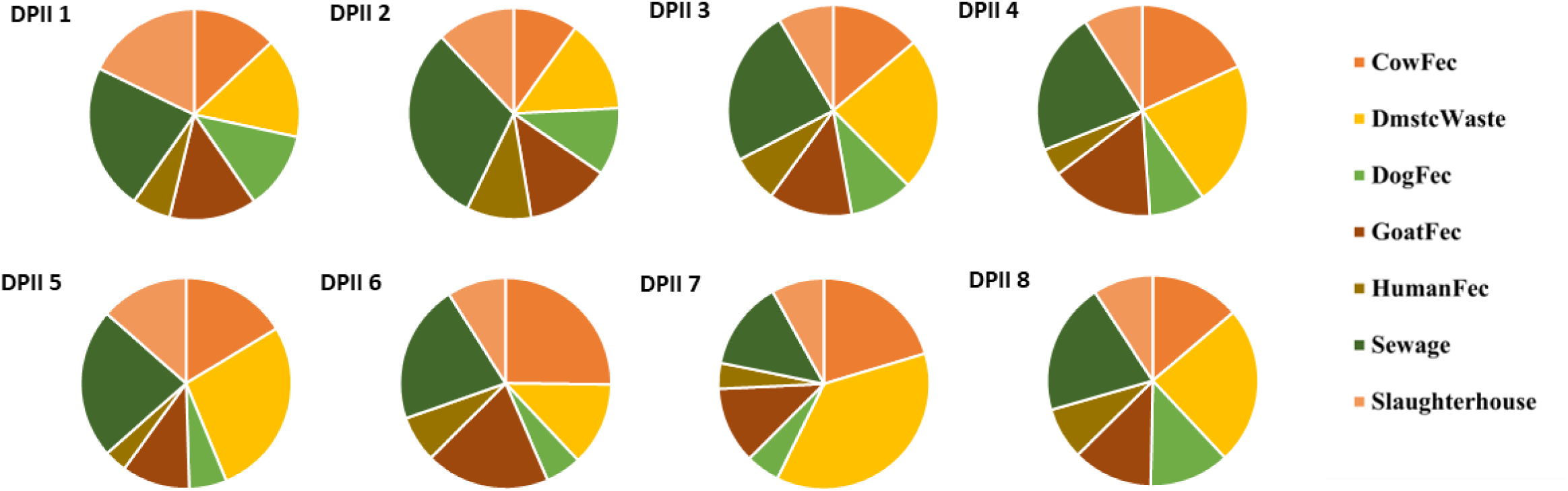
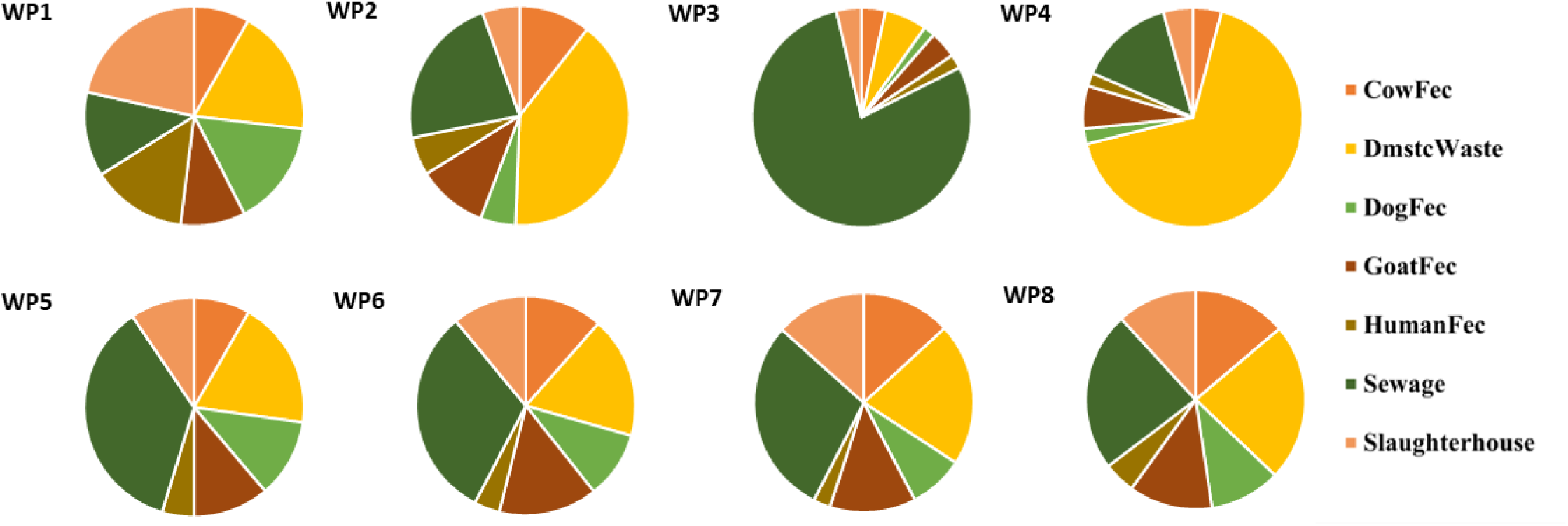
NMDS analysis of source and estuarine samples, where 1-8 denote the sampling Stations, WP is wet period (June-Sept), DP I is dry period I (Feb-May), and DP II is dry period II (Oct-Jan).

Similarly, the PCoA plots (**Fig. 6**) revealed that the sewage and domestic waste samples clustered more closely to the estuarine samples, compared to the animal fecal samples and slaughterhouse waste. The PCoA plot using unweighted UniFrac distance (plot (a) in **Fig. 6**) measures similarities between samples based on the presence or absence of bacterial species, whereas the plot using weighted UniFrac distance (plot (b) in **Fig. 6**), compares similarities between samples based on both presence/absence and abundance of bacterial species. Both plots illustrate how the seasons change the bacterial communities as well. During the post-monsoon months (S/DP I), Stations 4, 5, 6, and 7 appear to be the most dissimilar to all source samples. Station 6 appears to be polluted with slaughterhouse waste during DP II (winter months).

**Fig. 6.**
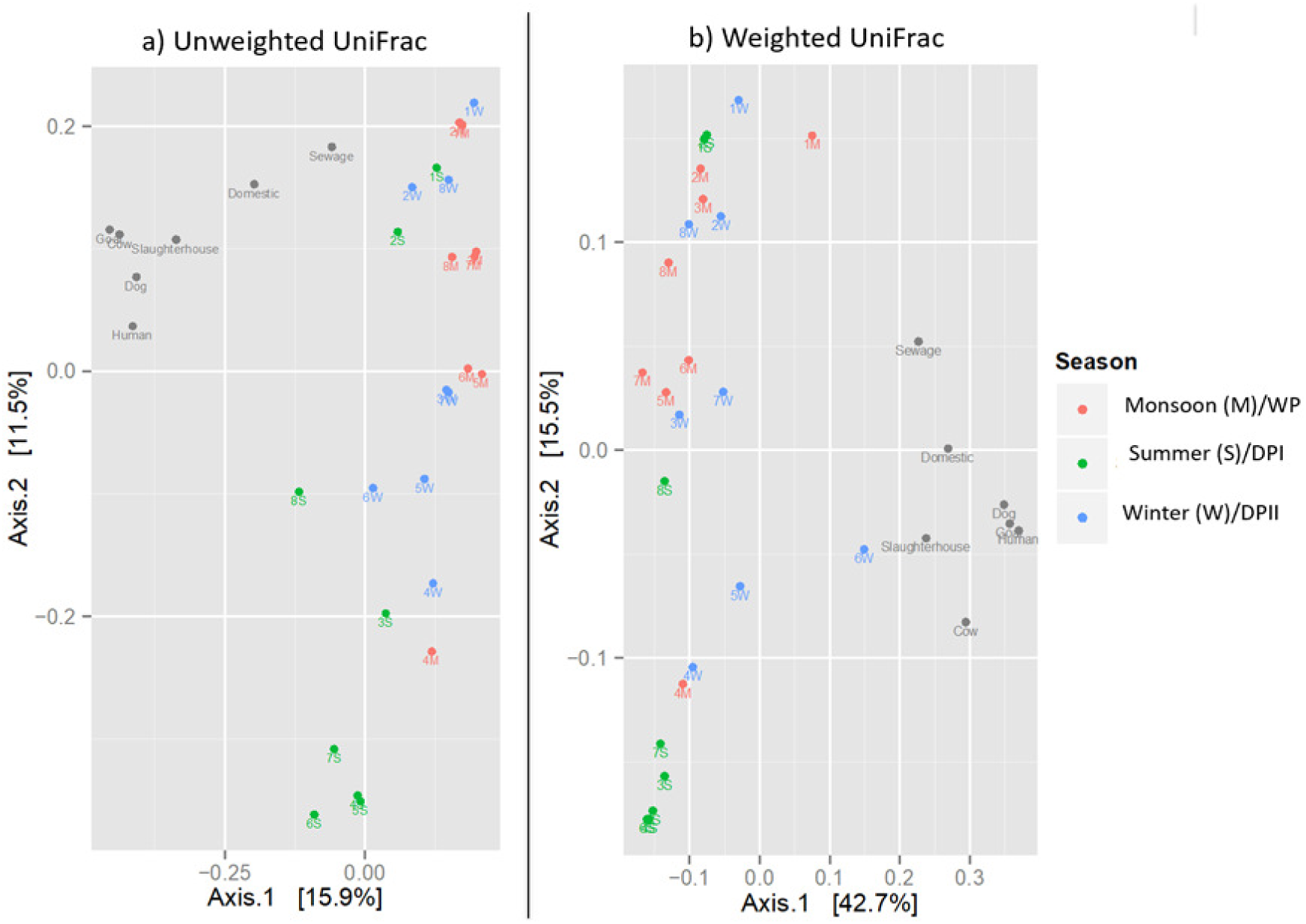
PCoA plots with a) unweighted and b) weighted UniFrac distance measurements

### Source Tracker analysis

One of the most potent and successful applications for interpreting NGS data to track microbial sources was the SourceTracker application, which indicated that the major microbial contaminants in the estuarine water samples were sewage and domestic waste in all seasons. The shared microbial communities comprised 45-94% of the estuarine samples and 0.18-32.76% of the source samples. In the wet period (WP), the environmental water samples were more contaminated by domestic waste and sewage. At Station 3, the percentage of sewage contamination was almost 79%. In other Stations, the percentage of sewage contamination ranged from 12-36% in the WP. Domestic waste constituted about 67% and 40% in Station 4 and Station 2 respectively. In other stations, the domestic waste was ranged from 6-23% (**Fig. 7a**). Similar to WP, in DP I and II, 50-60% of the microbial contaminants were from domestic waste and sewage, except in Station 6, where the major contaminant was human feces, which constituted about 60% in DP I. In DP II, domestic waste contributed 37% at Station 7 as well as 13-27% at all the other Stations. In DP I and DP II, the percentage of human feces contamination was less compared to sewage and domestic waste, ranging from 3-14% **(Fig. 7b, c).**

**Fig. 7.**
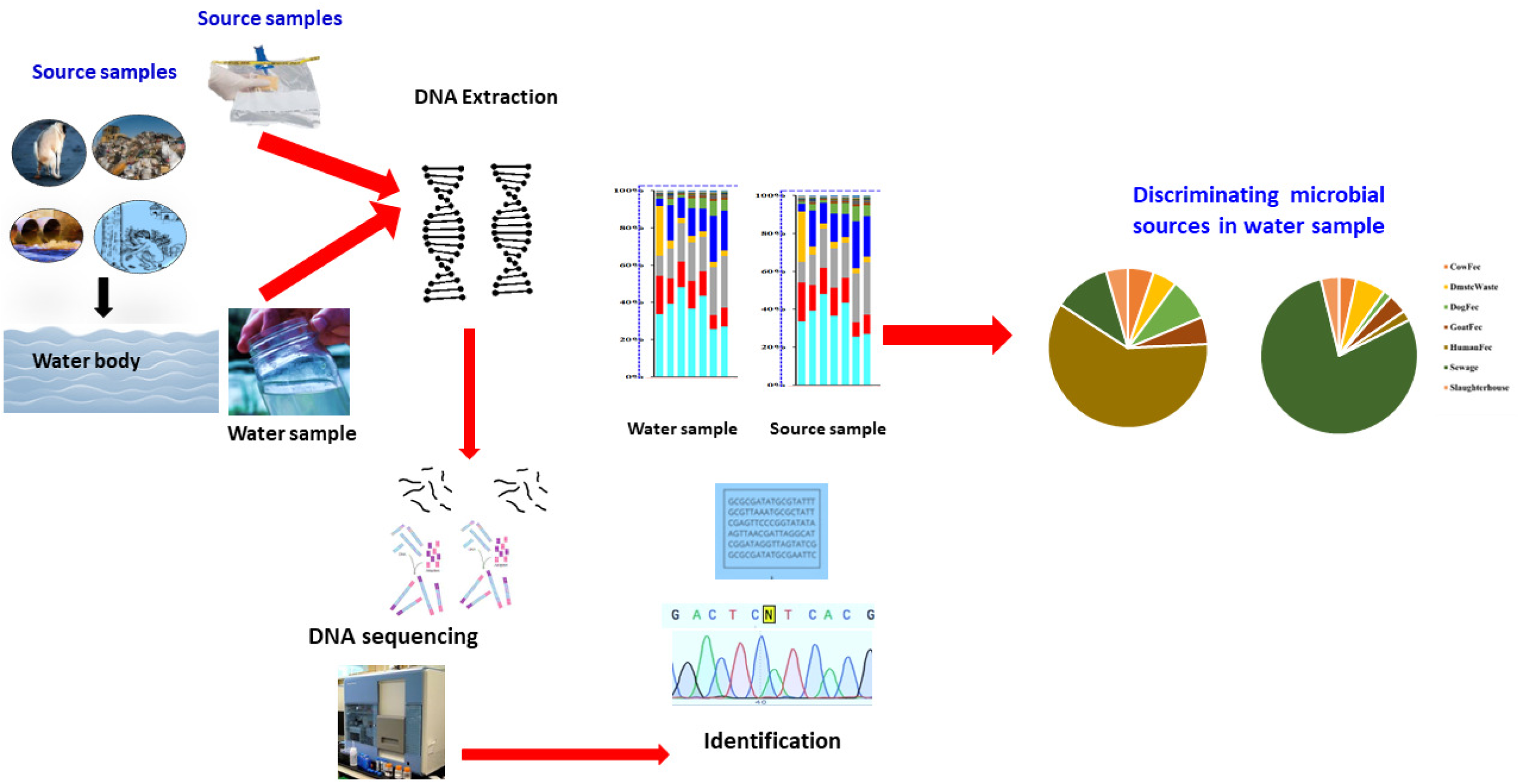
Pie charts representing source tracking of microbial contaminates from 8 stations in CE, where 1-8 denote the sampling Stations, WP is the wet period (June-Sept), DP I is dry period I (Feb-May), and DP II is dry period II (Oct-Jan).

In domestic waste and fecal samples, the dominant family was *Clostridiaceae*. They are commonly found in various environments, and some species can be found in the intestines of humans and other animals as part of the normal gut microbiome. In almost all the stations in the dry and wet periods, the *Clostridiaceae* was present (0.54 - 4.70%). Their abundance was comparatively higher in DP II, especially in the southern region **(Fig. 3)**. The family *Bacteroidaceae*, comprised of *Bacteroides,* are the most abundant and prevalent bacteria in the human gut, especially in the colon (Amini Khiabani et al., 2023). In DP I and II, *Bacteroidaceae* were found mostly at stations 2, 4, and 6, while they were present at Stations 1, 3, 5, and 7 during WP (**Fig. 3)**. Their abundance was relatively high (7.3%) at Station 6 during DP I. This Station also had high human fecal contamination (60%) (**Fig. 7)**.

The *Prevotellaceae* and *Ruminococcaceae* families are both important groups of bacteria found in the human gut microbiome and were detected in human feces in the present study. In DP I at Station 6, the *Prevotellaceae* family constituted about 13.8% and the *Ruminococcaceae* family constituted about 4% of the communities in the estuarine samples. The *Enterobacteriaceae* family present in the estuarine samples ranged from 0.09 to 1.44%, whereas in the source samples, it ranged from 0.12 to 3.56%. In the dry period, *Enterobacteriaceae* were most abundant at Station 2 (1.44%) and Station 8 (1.04%). The *Lachnospiraceae* are important butyrate producers residing in the intestinal microbiome, and they constituted 6% of DP I at Station 6 (**Fig. 7**). The *Rhodocyclaceae* family was majorly found in sewage and domestic waste and was present at almost all stations in wet and dry periods. The *Microbacteriaceae* family was present in cow, dog, and goat feces and was abundant in DP II, especially in the southern region of the CE **(Fig 3)***. The Flavobacteriaceae* family was present in all source samples with a higher percentage in domestic waste followed by dog feces and sewage samples. In all the estuarine samples, *Flavobacteriaceae* were present at higher percentages during DP II than other seasons.

## Discussion

The Cochin Estuary (CE), one of the largest tropical wetland habitats on India’s southwest coast located adjacent to the Arabian Sea, served as the site of the current study. According to Gupta et al., (25), the untreated dumping of industrial wastes (1.04×10^5^ m^3^d^−1^) and home sewage (260 m^3^d^−1^) into the CE has caused detrimental alterations in the ecosystem, including changes in the nutrient stoichiometry and even microbiological activity (26, 27). In India, sewage is not chemically treated and just 12.45% of sewage is properly handled before it is dumped onto the ground or into water bodies (28). In Kochi, there are only two operating sewage treatment plants (STPs), and they use activated sludge tanks to reduce the organic load in the primary effluents (29). Municipal sewage treatment facilities are thought to be “hotspots” for the release of antibiotic-resistant bacteria (ARB) and antibiotic-resistance genes (ARGs) into the aquatic environment (30, 31).

There are only limited studies on microbial communities in estuarine environments compared to water and sediments of coastal and deep-sea oceans, lakes, and sewage. Freshwater that is rich in nutrients is continuously pumped into the CE through riverine inputs (26,27). Additionally, because of its wide mouth, seawater from the Arabian Sea can enter the entire estuary, creating a distinct environmental gradient **(Fig. 1)**. There have been reports of seasonal and geographic variations in the ecological factors (abiotic or biotic) in the water and sediment in the CE (32). Additionally, due to rapid development, increased agricultural activity, and wastewater runoff, the CE and its surroundings experience severe anthropogenic pollution (26,27). The CE is a suitable area for assessing microbial diversity, responses to environmental instabilities, and community structure dynamics, as well as for testing ecological theories, given the variety of natural circumstances and human disturbances. In the present study, changes in estuarine hydrography and pollution caused variations in the relative abundance of bacterial communities in CE, despite being composed of aquatic microbial communities that are similar to those found in other aquatic environments (27, 33–35). In many ecosystems, salinity is a significant environmental element that shapes bacterial communities (27, 36–39).

Our results revealed that even though there is a clear variation in the physicochemical parameters, especially salinity, 90% of sequence reads at the phylum level were shared among the 8 sites sampled from the CE, especially the predominance of phylum Proteobacteria. Bacterioplankton community analysis based on Illumina sequencing from four Sundarbans estuaries, namely Mooriganga, Thakuran, Matla, and Harinbhanga, and also from other estuaries such as Columbia, Delaware, Jiulong, Pearl, and Hangzhou also revealed that Proteobacteria dominated all nine estuaries (40). Other tropical estuaries such as Pearl River Estuary (41), Cochin Estuary (27), Zuari Estuary (35, 42), and Serinhaém Estuary (43) also support the same result. According to these reports, Bacteroidetes, Firmicutes, Actinobacteria, Acidobacteria, Verrucomicrobia, Planctomycetes, and Cyanobacteria were other abundant phyla in these estuaries (40, 44, 45).

Diverse microbial communities have been found in wastewater in several studies, and these communities may have an impact on the microbial ecology of these ecosystems (46). Proteobacteria predominate in wastewater, according to recent research on the global diversity of bacterial communities in 269 wastewater treatment plants (47). In the present study, Proteobacteria were dominant in sewage samples (10-35%). This phylum is known to play a wide range of roles in the breakdown of organic matter and the cycling of nutrients (48). 80% of fecal microbiota are typically members of three phyla: Bacteroidetes, Firmicutes, and Actinobacteria (49, 50). In general, the Firmicutes to Bacteroidetes ratio is thought to be important in the composition of human and animal gut microbiomes (51). The bacterial families such as *Clostridiaceae, Bacteroidaceae*, and *Prevotellaceae* were detected in fecal samples based on MST (52).

Station 3 is located near a region with high population density, which leads to a greater volume of sewage generation, leading to sewage contamination of estuarine water. There are recent reports from the Kerala Water Authority (KWA) that water bodies and groundwater are becoming polluted due to sewage contamination (53, Times of India Report). The present study showed that Station 3 had 79% sewage contamination. Natural factors such as heavy rainfall in wet periods can overwhelm sewage systems, causing sewage overflow and contamination of nearby estuarine water bodies. Improper waste disposal, open defecation, and lack of awareness about sanitation practices can also contribute to sewage contamination. KWA is planning to connect around 8,000 houses in six divisions in the region near Station 3 to a proposed STP with a capacity of 9 million liters per day (MLD) to address sewage pollution issues in the area (54, Times of India Report). Our study results, indicating that this region is heavily contaminated by sewage, are correlated with this report and underscore the need for intervention. In Stations 2 (Periyar River) and 4 (Cochin Inlet from the Arabian Sea), the percentage of domestic waste was higher (40% and 67%, respectively) compared to other stations in the wet period. In these stations, the fecal contaminants from houseboats, garbage from nearby towns, and organic pollutants from nearby agriculture fields might be the predominant reasons for the contamination.Periyar River and its tributaries are entering these estuarine systems at S2 and S4 stations. Regardless of salinity, increased total coliform values suggest the presence and survival of enteric bacteria in this system (55).

High population density and rapid urbanization might have strained existing sanitation systems, leading to a higher volume of untreated waste being released into the environment. Inadequate STPs or the absence of proper sewage infrastructure may have resulted in untreated or partially treated sewage entering the estuary. Station 6 in the CE acts as a mesohaline zone in the DP I and is physiochemically less dynamic as a result of low flushing rates during dry seasons with no rainfall and reduced riverine inputs. Our study found that human fecal contamination was high at Station 6 compared to other pollution samples (**Fig. 7),** and high microbial activity has been reported previously (39). The relatively high human fecal contamination found at Station 6 may be influenced by the lack of flushing of organic contaminants due to the presence of a bund near Station 8 that prevents freshwater from Vembanad Lake from entering CE to protect paddy fields to the south, this will alter the estuary’s fresh and saltwater balance (39, 26).

The microbiological flora of soil, food, vegetables, and animal and human wastes make up the majority of domestic waste. Estuaries are the principal recipients of pollutants from point sources, such as hospitals, agricultural fields, and industrial and sewage outfalls, as well as non-point sources, such as rainwater runoff, ballast water, and animal feces (56–58). The water quality of CE is degraded by industrialization and urbanization in Cochin City and adjacent areas (56, 59), including sewage and residential waste. On the other hand, houseboat tourism (houseboats, private motorboats, and speed-boats) discharges wastewater of 0.23 million liters per day (https://timesofindia.indiatimes.com/city/kochi/vembanad-hotbedofsuperbugs/articleshow/56341387.cms). Significantly, the estuary and coastal waters accumulate pollution indicator species more in the wet period than the dry period, indicating that rainfall causes an exponential increase in bacterial biomass as well as increased turbidity in the water column (60). Understanding the seasonality of bacterial contamination will aid in management solutions.

## Materials and Methods

### Site description and sampling

The CE is a tropical estuarine system within the Vembanad Lake region, extending between *9°58′N latitude and 76°15′E longitude*. Its dimensions span 80 km in length and vary in width from 500 to 4000 m. The estuary maintains a depth range of 2 to 7 m, with shipping routes dredged to 10-13 m. The CE has two permanent inlets to the Arabian Sea at Kochi and Azhikode. Six major rivers - Pamba, Achancovil, Manimala, Meenachil, Periyar, and Muvattupuzha – Notably, the Periyar and Muvattupuzha Rivers contribute to the CE’s freshwater influx during the southwest monsoon, altering its salinity from 30 ppt at the entrance to 0.2 ppt at the river confluences (**Fig, 1**).

Water samples were collected from eight distinct sites along the Periyar River and Vembanad Lake during the monsoon (June-September), post-monsoon (October-January), and pre-monsoon (February-May) periods (**Fig. 1**). The sampling sites were selected to represent various aspects of the backwater system around Kochi, Kerala. The upstream sites (Stations 1 −3) reflect relatively unpolluted river stretches, while downstream sites (Stations 5-8) encompass industrial and coastal areas (**Fig. 1**).

### Environmental water samples

Water sample collection protocols involved using sterilized 1-L glass bottles to collect samples in triplicate at each site to mitigate variability and ensure data accuracy. The collected samples were immediately placed on ice and shielded from the sun while in the field. Environmental parameters, such as salinity and temperature, were measured using a portable CTD (SBE, Seabird Electronics, USA). Upon returning to the lab, the samples were filtered using 0.2-μm filters (Millipore, USA), and the filtered papers were stored at −80 °C until further processing (27).

### Environmental Parameters

At each sampling site, additional water samples were collected for analysis of nitrate (NO_2_), nitrite (NO_3_), phosphate (PO_4_), ammonia (NH_4_), silicate (SiO_4_), dissolved oxygen (DO), and pH. Samples were collected in clean 500 mL plastic bottles and immediately placed on ice. Upon returning to the lab, samples were frozen at −20 °C until analysis. Samples for nutrients (ammonia, nitrite, nitrate, phosphate, and silicate) were filtered and estimated spectrophotometrically by standard protocols (61). Dissolved oxygen (DO) content in the ambient water was determined following Winkler’s titration method (61).

### Bacterial source samples

Bacterial source samples included raw sewage, landfill leachate, household wastewater, slaughterhouse waste, and feces from humans, cows, dogs, and goats. Influent sewage was collected at various sewage treatment plants throughout the region. Household wastewater was collected from open roadside canals that also capture rainwater runoff within the city. Slaughterhouse waste was collected from poultry, beef, and goat slaughterhouses around Kochi near the backwaters. A total of 8 human feces samples were collected from hospitals in and around Cochin. Similarly, cow, dog, and goat fecal samples were collected fresh from various locations in and around Cochin and the Cochin backwaters region. All samples were collected in sterile WhirlPak (Nasco, Wisconsin, USA) sampling bags and immediately placed on ice and shielded from the sun. Upon returning to the lab, samples were frozen at −80°C until further processing.

### DNA isolation and sequencing

Under aseptic conditions, filter papers (i.e., Water samples were filtered using 0.2-μm filters-Millipore, USA) from the same location and season were cut into small pieces using sterilized scissors and mixed. Similarly, fecal samples from the same source were homogenized using sterilized stainless-steel spoons and bowls. The Power-Soil DNA Isolation Kit (Mo Bio Laboratories, California, USA) was used to extract total metagenomic DNA from the water and bacterial source samples following the manufacturer’s protocol with modifications (13). Slaughterhouse waste was homogenized using sterilized scissors, and its DNA was extracted using the Qiagen Blood and Tissue kit (Hilden, Germany). The quality of the DNA was checked using Nanodrop (Thermo Fisher Scientific, USA) and subsequently stored at −80°C. For library construction and next-generation sequencing, DNA was sent to GenePath Dx (Pune, India) (62).

### Amplicon library construction and Illumina sequencing

The library construction was initiated after the DNA quantification (minimum concentration of 10 ng/ul) using Broad Range Qubit System (Life Technologies, CA, USA). For sequencing PCR targeting a product size of 400-600bp using V3-V4region along with 20bp adapters with standard PCR conditions (95°C-3 min, 40 cycles of 95 °C-15 sec, 60 °C-20 sec 72 °C-10 min was used. Indexing PCR was performed with QuantiTect MultiPlex PCR kit (Qiagen, Germany) using index primers, adapters, and unique barcode sequences. The cycling conditions comprised of 95°C-15 minutes, succeeded by 30 cycles of 95°C-30 sec, 63°C-45 sec, and 72°C-90 sec. After PCR, subsequent quantification of the amplified products was performed in the Broad Range Qubit System (Life Technologies, CA, USA) (63).

The PCR products (minimum size of 300 bp) were purified with the PureLink PCR Purification kit (Invitrogen in California, USA. Then by using the Illumina library preparation kit, libraries were prepared for Next-Generation Sequencing (NGS). In an Illumina MiSeq v3 cartridge, prepared samples were loaded and sequencing was carried out in an Illumina MiSeq next-generation sequencer (2*300 mode) (63,64).

### Initial processing of sequence reads & statistical analysis

After sequencing, adapter sequences were removed. Preprocessing such as demultiplexing, quality filtering (scores above Q30), OTU picking, and taxonomic assignment (Greengenes database) was carried out on the Illumina BaseSpace Labs website (Version 1.0.0). The QIIME-based software such as TagCleaner software, bcl2fastq Conversion Software, the automated FASTQ Tool Kit application, etc were used for analysis (62, 64).

PERMANOVA and analysis of similarities (ANOSIM) were performed to understand significant variations in space and time. The species abundance was normalized and transformed using the Bray-Curtis method. The potential relationships between all microbial and environmental variables were tested using the Karl Persons Correlation Analysis. The similarity profile routine (SIMPROF) technique and non-metric multidimensional scaling (NMDS) were used to segregate or spatially group bacterial communities across seasons (PRIMER 6). Weighted and unweighted UniFrac distances were calculated in QIIME, and principal coordinate analysis (PCoA) was performed with the UniFrac distances using Phyloseq in R to visualize the similarities between the water and bacterial source samples. Clustering was performed using the Euclidean distance matrix and the group average approach. We utilized SourceTracker, a Bayesian algorithm, to assess the chance that an OTU found in one community (e.g., the environmental sample) could be from another community (e.g., the source) to identify the source of OTUs (https://github.com/danknights/sourcetracker; 65).

### Nucleotide sequence accession number

The data generated in this study was deposited in NCBI Sequence Read Archive with SRA: PRJNA595444

## Acknowledgments

The authors thank the Director, NIO, Goa, and the Scientist-in-Charge, CSIR-NIO (RC), Kochi for their support and advice. This study was part of the *Fulbright Fellowship* (USIEF) of MC. JV is grateful to the Council of Scientific and Industrial Research (CSIR), New Delhi for the Senior Research Fellowship Grant (award letter 31/53 (0037)/2014-EMRI) and the UGC-Kothari postdoctoral fellowship (BL/20-21/0310 (S-90). *JV and AP are* also thankful to the Department of Marine Biology, Microbiology and Biochemistry, and the Department of Biotechnology, Cochin University of Science and Technology. This is NIO contribution number xxxx.

## Authors Contributions

AP: Conceptualization, Supervision, Investigation, Writing - review & editing, and Funding acquisition

JV: Methodology, Formal analysis, Investigation, Data curation, Writing - original draft

MC: Conceptualization, Methodology, Funding acquisition, Formal analysis, Investigation, Writing - review & editing

DGP: Statistical Analysis

NP: Next generation sequencing

MN: Chemical Analysis

## Author Statement

We confirm that: The manuscript has been read and approved by all co-authors. The work complies with ethical standards and, where applicable, has received approval from relevant institutional or ethical committees. There are no conflicts of interest related to this research, or any potential conflicts have been disclosed. All sources of funding and support have been acknowledged. Data supporting the findings of this study are available upon reasonable request.

## Data availability

Data will be made available on request.

## Consent to participate

Not applicable.

## Consent for publication

The manuscript must have written consent from all the authors and from the institution.

## Supplementary Materials

**Fig. S1** Class level distribution of microbial communities, where 1-8 denote the sampling Stations, WP is wet period (June-Sept), DP I is dry period I (Feb-May), and DP II is dry period II (Oct-Jan).

**Table S1** Environmental water parameters, where 1-8 denote the sampling Stations, WP is wet period (June-Sept), DP I is dry period I (Feb-May), and DP II is dry period II (Oct-Jan).

## References

1. Jalloul, G., Keniar, I., Tehrani, A., & Boyadjian, C. (2021). Antibiotics Contaminated Irrigation Water: An Overview on Its Impact on Edible Crops and Visible Light Active Titania as Potential Photocatalysts for Irrigation Water Treatment. Frontiers in Environmental Science, 9, 767963. 10.3389/fenvs.2021.767963

2. Shayo, S., et al. (2023). Diversity of waterborne pathogens in aquatic environments: A global review. Environmental Research, 214, 113923. 10.1016/j.envres.2023.113923

3. Valério, E., Santos, M. L., Teixeira, P., Matias, R., Mendonça, J., Ahmed, W., & Brandão, J. (2022). Microbial source tracking as a method of determination of beach sand contamination. International Journal of Environmental Research and Public Health, 19(13), 7934.doi: 10.3390/ijerph19137934

4. Harwood, V.J., Staley, C., Badgley, B.D., Borges, K., & Korajkic, A. (2014). Microbial source tracking markers for detection of fecal contamination in environmental waters: relationships between pathogens and human health outcomes. FEMS Microbiology Reviews, 38(1), 1–40. 10.1111/1574-6976.12031

5. Corrigan, J. A., Butkus, S. R., Ferris, M. E., & Roberts, J. C. (2021). Microbial source tracking approach to investigate fecal waste at the strawberry creek watershed and clam beach, California, USA. International Journal of Environmental Research and Public Health, 18(13), 6901. 10.3390/ijerph18136901

6. Murugan, A., et al. (2011). Influence of anthropogenic activities on microbial diversity in tropical estuaries. Marine Pollution Bulletin, 62, 1352–1360. 10.1016/j.marpolbul.2011.04.019

7. Holcomb, D. A., & Stewart, J. R. (2020). Microbial indicators of fecal pollution: recent progress and challenges in assessing water quality. Current Environmental Health Reports, 7, 311–324. 10.1007/s40572-020-00278-1

8. Li, E., Saleem, F., Edge, T. A., & Schellhorn, H. E. (2021). Biological indicators for fecal pollution detection and source tracking: A review. Processes, 9(11), 2058. 10.3390/pr9112058

9. González-Fernández, A., Symonds, E. M., Gallard-Gongora, J. F., Mull, B., Lukasik, J. O., Navarro, P. R., & Harwood, V. J. (2021). Relationships among microbial indicators of fecal pollution, microbial source tracking markers, and pathogens in Costa Rican coastal waters. Water Research, 188, 116507. 10.1016/j.watres.2020.116507

10. KSPCB IN OA NO. 639 (2019) National Green Tribunal https://greentribunal.gov.in › news_updates › R… PDF On 14/11/2019 the Kerala State Pollution Control Board rules.2019 https://nmcg.nic.in/writereaddata/fileupload/ngtmpr/0Kerala%20MPR%2October%202020.pdf

11. Shankar, A., Jagajeedas, D., Radhakrishnan, M. P., Paul, M., Narendrakumar, L., Suryaletha, K., … &Thomas, S. (2021). Elucidation of health risks using metataxonomic and antibiotic resistance profiles of microbes in flood affected waterbodies, Kerala 2018. Journal of Flood Risk Management, 14(1), e12673. 10.1111/jfr3.12673

12. Padua, S., Kripa, V., Prema, D., Mohamed, K. S., Jeyabaskaran, R., Kaladharan, P., … & Babu, A. (2023). Assessment of ecosystem health of a micro-level Ramsar coastal zone in the Vembanad Lake, Kerala, India. Environmental Monitoring and Assessment, 195(1), 95. 10.1007/s10661-022-10692-7

13. Cao, Y., Van De Werfhorst, L.C., Dubinsky, E.A., Badgley, B.D., Sadowsky, M.J., Andersen, G.L., Griffith, J.F., Holden, P.A. 2013. Evaluation of molecular community analysis methods for discerning fecal sources and human waste. Water Research. 47: 6862–6872. 10.1016/j.watres.2013.02.061

14. Griffith, J.F., Layton, B.A., Boehm, A.B., Holden, P.A., Jay, J.A., Hagedorn, C., McGee, C.D., Weisberg, S.B. 2013. The California Microbial Source Identification Manual: A Tiered Appraoched to Identifying Fecal Pollution Sources to Beaches. Southern California Coastal Water Research Project, Technical Report 804. Retrieved from:ftp://ftp.sccwrp.org/pub/download/DOCUMENTS/TechnicalReports/804_SIPP_MST_ManualPag.pdf

15. McLellan S.L., Huse S.M., Mueller-Spitz S.R., Andreishcheva EN, Sogin ML. 2010. Diversity and population structure of sewage-derived microorganisms in wastewater treatment plant influent. Environmental Microbiology. 12(2): 378 –392. DOI: 10.1111/j.1462-2920.2009.02075.x

16. Unno, T., Jang, J., Han, D., Kim, J.H., Sadowsky, M.J., Kim, O.-S., Chun, J., Hur, H.-G., 2010. Use of barcoded pyrosequencing and shared OTUs to determine sources of fecal bacteria in watersheds. Environmental Science and Technology. 44(20): 7777–7782. DOI: 10.1021/es101500z

17. Jeong J.Y., Park H.D., Lee K.H., Weon H.Y., Ka J.O. 2011. Microbial community analysis and identification of alternative host-specific fecal indicators in fecal and river water samples using pyrosequencing. Journal of Microbiology. 49(4): 585–594. DOI: 10.1007/s12275-011-0530-6

18. Lee, J.E., Sunghee, L., Sung, J., GwangPyo, K. 2011. Analysis of human and animal fecal microbiota for microbial source tracking. The ISME Journal. 5:362–365. doi: 10.1038/ismej.2010.120

19. Newton R.J., Bootsma M.J., Morrison H.G., Sogin M.L., McLellan S.L. 2013. A microbial signature approach to identify fecal pollution in the waters off an urbanized coast of Lake Michigan. Microbial Ecology. 65(4): 1011–1023. DOI: 10.1007/s00248-013-0200-9

20. Neave, M., Luter, H., Padovan, A., Townsend, S., Schobben, X., & Gibb, K. (2014). Multiple approaches to microbial source tracking in tropical northern Australia. MicrobiologyOpen, 3(6), 860–874.10.1002/mbo3.209

21. Ahmed, W., Staley, C., Sadowsky, M.J., Gyawali, P., Sidhu, J.P.S., Palmer, A., Beale, D.J., Toze, S. 2015. Toolbox approaches using molecular marker and 16S rRNA gene amplicon data sets for identification of fecal pollution in surface water. Applied and Environmental Microbiology. 81(20): 7067–7077. DOI: 10.1128/AEM.02032-15

22. Vierheilig, J., Savio, D., Ley, R. E., Mach, R. L., Farnleitner, A. H., & Reischer, G. H. (2015). Potential applications of next generation DNA sequencing of 16S rRNA gene amplicons in microbial water quality monitoring. Water Science and Technology, 72(11), 1962–1972.Bibbal, D., Um, M. M., Diallo, A. A., Kérourédan, M., Dupouy, V., Toutain, P. L., … & Brugère, H. (2018). Mixing of Shiga toxin-producing and enteropathogenic *Escherichia coli* in a wastewater treatment plant receiving city and slaughterhouse wastewater. International Journal of Hygiene and Environmental Health, 221(2), 355-363. 10.1016/j.ijheh.2017.12.009

23. Boukerb, A. M., Noël, C., Quenot, E., Cadiou, B., Chevé, J., Quintric, L., … & Gourmelon, M. (2021). Comparative analysis of fecal microbiomes from wild waterbirds to poultry, cattle, pigs, and wastewater treatment plants for a microbial source tracking approach. Frontiers in Microbiology, 12, 697553. 10.3389/fmicb.2021.697553

24. Zhao, X. L., Qi, Z., Huang, H., Tu, J., Song, X. J., Qi, K. Z., & Shao, Y. (2022). Coexistence of antibiotic resistance genes, fecal bacteria, and potential pathogens in anthropogenically impacted water. Environmental Science and Pollution Research, 29(31), 46977–46990. 10.1007/s11356-022-19175-1.

25. Gupta, et al., (2016)

26. John, S., Muraleedharan, K. R., Revichandran, C., Azeez, S. A., Seena, G., & Cazenave, P. W. (2020). What controls the flushing efficiency and particle transport pathways in a tropical estuary? Cochin Estuary, Southwest Coast of India. Water, 12(3), 908. 10.3390/w12030908.

27. Parvathi, A., Catena, M., Jasna, V., Phadke, N., & Gogate, N. (2021). Influence of hydrological factors on bacterial community structure in a tropical monsoonal estuary in India. Environmental Science and Pollution Research, 28(36), 50579–50592. 10.1007/s11356-021-14263-0

28. Central Pollution Control Board (CPCB) The Annual Report 2019-2020 https://cpcb.nic.in/annual-report.php

29. Central Pollution Control Board (CPCB) The Annual Report 2020-2021 https://cpcb.nic.in/annual-report.php

30. Um, M. M., Barraud, O., Kérourédan, M., Gaschet, M., Stalder, T., Oswald, E., … & Bibbal, D. (2016). Comparison of the incidence of pathogenic and antibiotic-resistant Escherichia coli strains in adult cattle and veal calf slaughterhouse effluents highlighted different risks for public health. Water research, 88, 30–38. DOI: 10.1016/j.watres.2015.09.029

31. Bibbal, D., Um, M. M., Diallo, A. A., Kérourédan, M., Dupouy, V., Toutain, P. L., … & Brugère, H. (2018). Mixing of Shiga toxin-producing and enteropathogenic Escherichia coli in a wastewater treatment plant receiving city and slaughterhouse wastewater. International Journal of Hygiene and Environmental Health, 221(2), 355–363. 10.1016/j.ijheh.2017.12.009.

32. Parvathi, A., Jasna, Vijayan., Jina, S., Jayalakshmy, K. V., Lallu, K. R., Madhu, N. V., and Balachandran, K. K (2015). Effects of hydrography on the distribution of bacteria and virus in Cochin estuary, India. Ecological Research (ISSN:1440-1703), 30(1), 85–92. 10.1007/s11284-014-1214-6.

33. Savio, D., Sinclair, L., Ijaz, U. Z., Parajka, J., Reischer, G. H., Stadler, P., … & Eiler, A. (2015). Bacterial diversity along a 2600 km river continuum. Environmental microbiology, 17(12), 4994–5007. 10.1111/1462-2920.12886

34. Meziti, A., Tsementzi, D., Ar. Kormas, K., Karayanni, H., & Konstantinidis, K. T. (2016). Anthropogenic effects on bacterial diversity and function along a river-to-estuary gradient in Northwest Greece revealed by metagenomics. Environmental Microbiology, 18(12), 4640–4652. 10.1111/1462-2920.13303

35. Eswaran R, Khandeparker L (2019) Seasonal variation in β-glucosidaseproducing culturable bacterial diversity in a monsoon-influenced tropical estuary. Environ Monit Assess 191(11):1–11. 10.1007/s10661-019-7818-0.

36. Liu, J., Fu, B., Yang, H., Zhao, M., He, B., & Zhang, X. H. (2015). Phylogenetic shifts of bacterioplankton community composition along the Pearl Estuary: the potential impact of hypoxia and nutrients. Frontiers in microbiology, 6, 64. 10.3389/fmicb.2015.00064

37. Liu, S. M., Qi, X. H., Li, X., Ye, H. R., Wu, Y., Ren, J. L., … & Xu, W. Y. (2016). Nutrient dynamics from the changjiang (yangtze river) estuary to the east china sea. Journal of Marine Systems, 154, 15–27. 10.1016/j.jmarsys.2015.05.010

38. Herlemann, D. P., Lundin, D., Andersson, A. F., Labrenz, M., & Jürgens, K. (2016). Phylogenetic signals of salinity and season in bacterial community composition across the salinity gradient of the Baltic Sea. Frontiers in Microbiology, 7, 1883. 10.3389/fmicb.2016.01883.

39. Jasna, V., Parvathi, A., Pradeep Ram, A. S., Balachandran, K. K., Madhu, N. V., Nair, M., … & Sime-Ngando, T. (2017). Viral-induced mortality of prokaryotes in a tropical monsoonal estuary. Frontiers in Microbiology, 8, 895. 10.3389/fmicb.2017.00895

40. Ghosh A, Bhadury P (2019) Exploring biogeographic patterns ofbacterioplankton communities across global estuaries. Microbiol Open 8(5):e00741. 10.1002/mbo3.741

41. Fan, Y., Li, Z., Li, B., Ke, B., Zhao, W., Lu, P., … & Kan, B. (2023). Metagenomic profiles of planktonic bacteria and resistome along a salinity gradient in the Pearl River Estuary, South China. Science of The Total Environment, 889, 164265. 10.1016/j.scitotenv.2023.164265.

42. Fernandes, V., & Bogati, K. (2019). Persistence of fecal indicator bacteria associated with zooplankton in a tropical estuary—west coast of India. Environmental monitoring and assessment, 191, 1–22. 10.1007/s10661-019-7531-z

43. De Santana, C. O., Spealman, P., Melo, V., Gresham, D., de Jesus, T., Oliveira, E., & Chinalia, F. A. (2021). Large-scale differences in diversity and functional adaptations of prokaryotic communities from conserved and anthropogenically impacted mangrove sediments in a tropical estuary. PeerJ, 9, e12229. 10.7717/peerj.12229.

44. Parvathi A, Jasna V, Aswathy VK, Nathan VK, Aparna S, Balachandran KK (2019) Microbial diversity in a coastal environment with coexisting upwelling and mud-banks along the south west coast of India. Mol Biol Rep 46:3113–3127. 10.1007/s11033-019-04766-y.

45. Sachithanandam V, Saravanane N, Chandrasekar K, Karthick P, Lalitha P, Elangovan SS, Sudhakar M (2020) Microbial diversity from the continental shelf regions of the Eastern Arabian Sea: a metagenomic approach. Saudi journal boil. Scie 27(8):2065–2075. 10.1016/j.sjbs.2020.06.011.

46. Giwa, A. S., Ali, N., Athar, M. A., & Wang, K. (2020). Dissecting microbial community structure in sewage treatment plant for pathogens’ detection using metagenomic sequencing technology. Archives of microbiology, 202, 825–833. 10.1007/s00203-019-01793-y

47. Wu, L., Ning, D., Zhang, B., Li, Y., Zhang, P., Shan, X., … & Zhou, J. (2019). Global diversity and biogeography of bacterial communities in wastewater treatment plants. Nature microbiology, 4(7), 1183–1195. 10.1038/s41564-019-0426-5

48. Nascimento, A. L., Souza, A. J., Andrade, P. A. M., Andreote, F. D., Coscione, A. R., Oliveira, F. C., & Regitano, J. B. (2018). Sewage sludge microbial structures and relations to their sources, treatments, and chemical attributes. Frontiers in Microbiology, 9, 1462. 10.3389/fmicb.2018.01462.

49. Lay C, Sutren M, Rochet V, Saunier K, Doré J, Rigottier-Gois L: Design and validation of 16S rDNA probes to enumerate members of the *Clostridium leptum* subgroup in human faecal microbiota. Environ Microbiol. 2005, 7: 933–946. 10.1111/j.1462-2920.2005.00763.x.

50. Li, C., Quan, Q., Gan, Y., Dong, J., Fang, J., Wang, L., & Liu, J. (2020). Effects of heavy metals on microbial communities in sediments and establishment of bioindicators based on microbial taxa and function for environmental monitoring and management. Science of the Total Environment, 749, 141555.10.1016/j.scitotenv.2020.141555

51. Li, W., & Ma, Z. (2020). FBA ecological guild: Trio of firmicutes-bacteroidetes alliance against actinobacteria in human oral microbiome. Scientific Reports, 10(1), 287. 10.1038/s41598-019-56561-1

52. Pantha, K., Acharya, K., Mohapatra, S., Khanal, S., Amatya, N., Ospina-Betancourth, C., … & Werner, D. (2021). Faecal pollution source tracking in the holy Bagmati River by portable 16S rRNA gene sequencing. NPJ Clean Water, 4(1), 12. 10.1038/s41545-021-00099-1.

53. Sreemol, TC. 2023. Soon, single network to transport sewage waste in Kerala. The Times of India. Accessed at: https://timesofindia.indiatimes.com/city/kochi/soon-single-network-to-transport-sewage-waste/articleshow/97851718.cms

54. Sreemol, TC. 2021. Kochi: KWA sewerage wing moots 7 decentralized STPs for corporation. The Times of India. https://timesofindia.indiatimes.com/city/kochi/kwa-sewerage-wing-moots-7-decentralized-stps-for-corp/articleshow/80352364.cms

55. TOI (Times of India). 2017. Vembanad, hotbed of superbugs. Time of India. Accessed at https://timesofindia.indiatimes.com/city/kochi/vembanad-hotbed-ofsuperbugs/articleshow/56341387.cms

56. Sukumaran D. P and Hatha A.M (2019) Screening of tropical estuarine water in south-west coast of India reveals emergence of ARGs-harboring hypervirulent Escherichia coli of global significance. International Journal of Hygiene and Environmental Health, 222(2), 235–248. 10.1016/j.ijheh.2018.11.002.

57. Guo, J., Li, J., Chen, H., Bond, P. L., & Yuan, Z. (2017). Metagenomic analysis reveals wastewater treatment plants as hotspots of antibiotic resistance genes and mobile genetic elements. Water research, 123, 468–478. 10.1016/j.watres.2017.07.002

58. Tanaka, Y., Minggat, E., & Roseli, W. (2021). The impact of tropical land-use change on downstream riverine and estuarine water properties and biogeochemical cycles. 10.1186/s13717-021-00315-3.

59. Salas, P. M., Sujatha, C. H., Kumar, C. R., & Cheriyan, E. (2017). Heavy metal distribution and contamination status in the sedimentary environment of Cochin estuary. Marine Pollution Bulletin, 119(2), 191–203.10.1016/j.marpolbul.2017.04.018.

60. Oliveira, D. B. C. D., Soares, W. D. A., & Holanda, M. A. C. R. D. (2020). Effects of rainwater intrusion on an activated sludge sewer treatment system. Revista Ambiente & Água, 15. 10.4136/ambi-agua.2497.

61. Grasshoff K (1999) Determination of nitrite, nitrate, oxygen, thiosulphate. In: Grasshoff K, Ehrhardt M, Kremling K (eds) Methods of seawater analysis. VerlagChemieWeinheim, NewYork, pp 139–142, 143–150, 61–72, 81–139–142, 143–150,61–72, 84.

62. Hill, P., Dextraze, M. F., Kroetsch, D., & Boddy, C. N. (2022). A comparison of hard and soft direct methods for DNA extraction from soil. bioRxiv, 2022-03. doi: 10.1101/2022.03.07.483395

63. Fernando, C., & Hill, J. E. (2023). cpn60 metagenomic amplicon library preparation for the Illumina Miseq platform. 10.21203/rs.3.pex-1438/v2.

64. Hu, T., Chitnis, N., Monos, D., & Dinh, A. (2021). Next-generation sequencing technologies: An overview. Human Immunology, 82(11), 801–811. 10.1016/j.humimm.2021.02.012

65. Du, B., Meng, L., Wu, H., Yang, H., Liu, H., Zheng, N., … & Wang, J. (2022). Source Tracker Modeling Based on 16S rDNA Sequencing and Analysis of Microbial Contamination Sources for Pasteurized Milk. Frontiers in Nutrition, 9, 845150. 10.3389/fnut.2022.845150.

